# Machine Learning Reveals The Effect of Maternal Age on The Mouse Pre-Implantation Embryo Developmental Timing

**DOI:** 10.1101/2022.05.17.492244

**Authors:** Nati Daniel, Tanya Wasserman, Zohar Adler, Tomer Czyzewski, Yonatan Savir

**Author notes:** These authors contributed equally to this work (Nati Daniel, Tanya Wasserman).

## Abstract

In recent years, many women have delayed childbearing, thus increasing the necessity for assisted reproductive technology (ART) for older women^1–3^. Despite advances in ART^4^, its success rate in advanced-age women is still very low^3,4^. As time-lapse imaging became available, morphological features of the developing pre-implantation embryo, *in-vitro*, are heavily used to assess its potency^5–9^. Timing of embryo cleavage is also an important factor that correlates with blastocyst formation and pregnancy rates^8,10–14^. Yet, our understanding of the interplay between embryos’ morphology, viability, and maternal age is limited, as manual approaches to infer embryo morphokinetics are time-consuming, subjective, and prone to errors. Machine learning^15–18^ was recently harnessed to predict embryo developmental potential^19,20^, however, with limited success.

Here, we develop an artificial intelligence (AI) platform that infers the embryos’ developmental stage and captures tens of morphological properties and developmental dynamics. We show that developmental timing is the most informative and predictive morphokinetic property, particularly for embryos from maternally aged females. Analyzing the timing distributions reveals that viable embryos are confined into an age-independent temporal corridor while non-viable embryos deviate from it towards slower transition times. Yet, the deviation of non-viable embryos from the temporal corridor is age-dependent. Furthermore, there is a significant correlation between consecutive developmental stages transition times that diminishes in maternally old embryos. Overall, our results suggest that maternally old embryos’ most apparent morphokinetic property is the loss of temporal regulation. Our results and platform pave the way for a more accurate, maternally-age-dependent, assisted reproductive technology.

## Results

Young (8-10 weeks) and maternally old (26-40 weeks) mouse females were induced to superovulate and mated with males. Eggs were then extracted and incubated in vitro (see Methods). The development of the embryos was monitored using a time-lapse microscope, from the zygote stage up to the late blastocyst stage. First, we curated a dataset to train and validate our pipeline. We annotated the different developmental stages in 3763 images of 89 embryos (from zygote to late blastocyst). 70% of the images were used as a training set, while the rest of the data was used for validation. Figure 1B illustrates the variability within similar developmental stages and demonstrates the challenge of identifying the developmental stage automatically.

**Figure 1.**
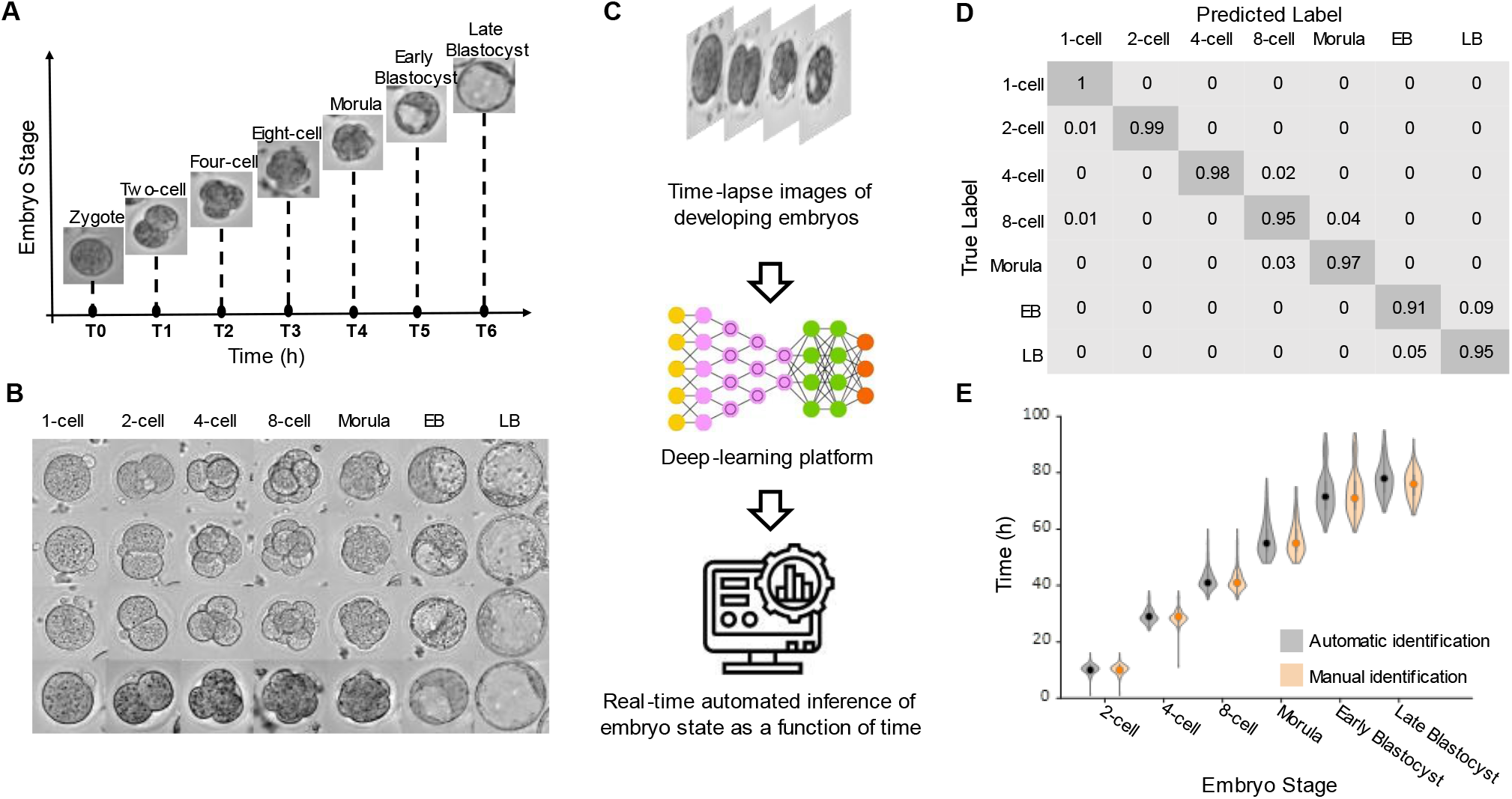
Machine learning platform automatically identifies the mouse pre-implantation developmental stages and their morphokinetic properties. A. Schematic overview of the developmental stages in pre-implantation mammalian embryo and the automatically inferred times by the deep learning platform of each corresponding developmental stage. B. An example from the created training set. C. Cropped consecutive images are fed to the developed deep learning platform. After the training, the developed platform is used to automatically identify the embryo stage in real-time. In addition, times and morpho-kinetic parameters are used to predict the blastocyst developmental potential. D. A confusion matrix of the developed AI platform in terms of developmental stage prediction (values are rounded). True positive rate (TPR) for each stage presented in the diagonal. E. Violin plot juxtaposing automatically- and manually-identified developmental times. EB denotes early blastocyst. LB denotes late blastocyst.

The pipeline we developed gets a video of the developing embryos as an input. The platform detects and tracks every single embryo during the entire video. The classification pipeline is composed of a pre-processing step for each embryo at each time point. This is followed by a deep convolutional network that is trained to classify the developmental stages according to the cropped embryo images (Methods). In addition, for each time point, the platform measures 45 morphological features of the embryo, including geometrical features (area, circumference, etc.), textural features (smoothness, spatial frequencies) (Methods, Figure 1C, Figure S1). We have tested multiple deep convolutional networks. ResNet-50 emerged as the best architecture for classifying the embryos’ developmental stages (Figure 1D, Figure S1).

Our results have state-of-the-art performance stage classification with an accuracy of 97.17%, macro-recall of 96.36% and macro-precision of 96.51%. This classification allows us to automatically infer, in real-time, the transition between developmental stages. To assess the ability of our platform to accurately measure the developmental transition times, we manually annotated the transition times of these 89 embryos. The mean difference between the manual and the automatic inferences for all the stages besides the late blastocyst stage is less than an hour. In the case of the late blastocyst stage, the differences are about two hours (Figure 1E). This stems from the fact that the definition of the late blastocyst stages involves estimating the ratio between the areas of the blastocoel and the embryo, which is challenging to do manually. To characterize the effect of maternal age on the viability and morphokinetic features of the developing embryo, we have conducted experiments on a second cohort of 103 embryos, 62 from young females and 41 from maternally old ones. We used a hybrid approach in which we verified the transition times within the time frame that the machine predicted. This allows both rapid and curated measurement of developmental timing.

Embryos from maternally aged females have a smaller probability of reaching the blastocyst stage. Figure 2A illustrates the survival curves of embryos from young and maternally old females. The fractions of embryos that reached the late blastocyst stage are 74% and 66% for embryos from young and old females, respectively. The main difference due to maternal age is the transition between the morula and early blastocyst stage (Figure 2A). An insightful way to probe the interplay between maternal age and the various morphokinetic features is to quantify the effect of knowing a given parameter on our estimation of the embryo survival probability. For example, if some feature has no impact on the survival probability, the added information of knowing it would be zero. We calculated the mutual information of each of the 46 features for the different stages and different ages (Figure 2B, Methods). Our results indicate that eight features are significantly informative in more than four stages, but the only one that is such for embryos from both maternally young and old females is developmental timing. Moreover, developmental timing is more informative in embryos from maternally old females (Figure 2B).

**Figure 2.**
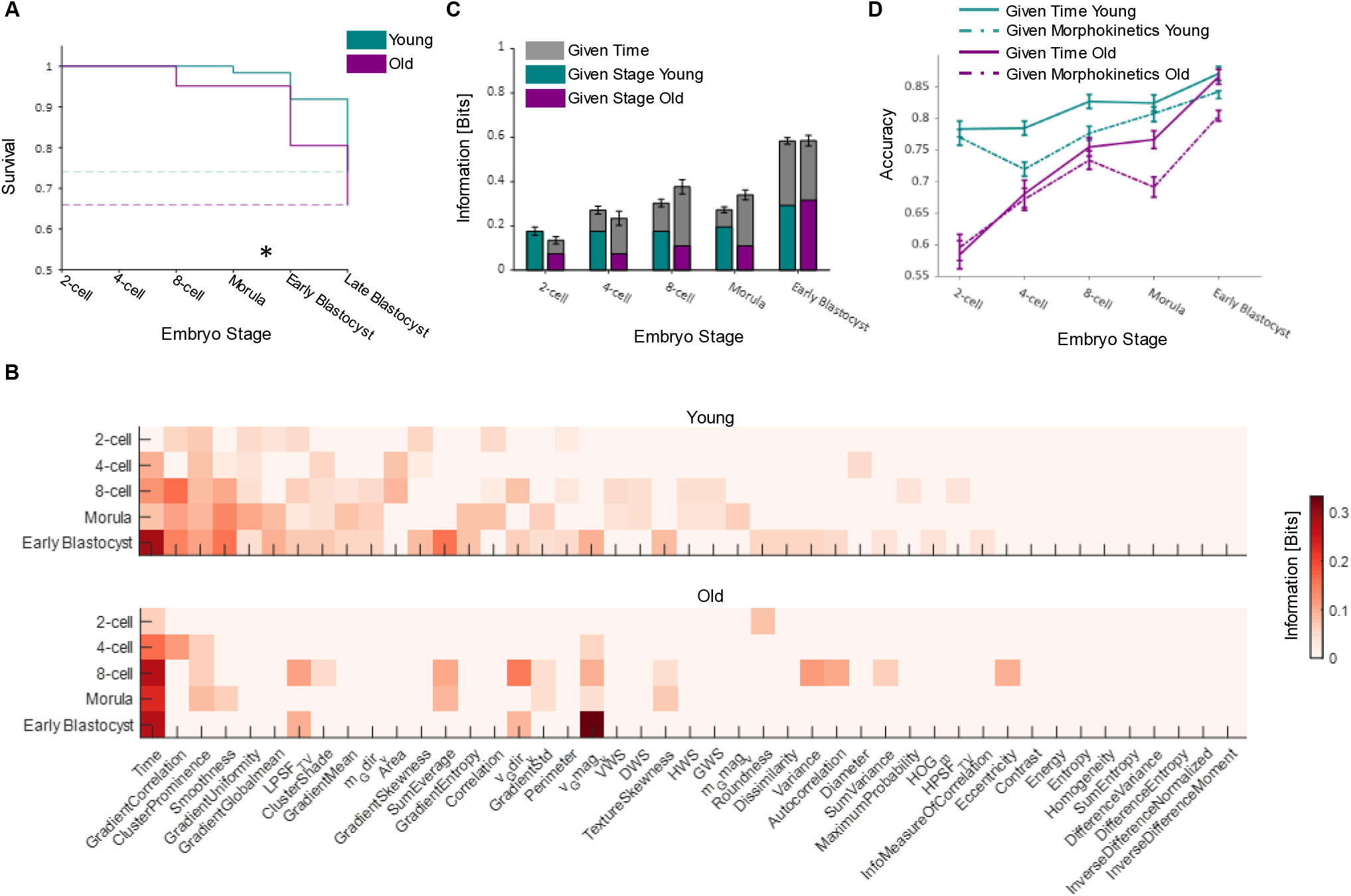
Timing is essential in predicting blastocyst formation rates, particularly in embryos from maternally aged females. A. Survival curves of *in-vitro* developing pre-implantation embryos derived from young and aged female mice (Young embryos n=62, Aged embryos n=41). Asterisk denotes significantly different stages (Binomial test, p-value<0.03). B. A heat map presenting the significant information extracted from each morphological parameter (total of 46 parameters) per developmental stage for maternally-young (upper panel) and aged (lower panel) embryos. The features are sorted according to the number of stages they are informative in C, for maternally-young embryos. Bar plot comparison between maternally young and aged embryos of the extracted information given the developmental stage with and without time information. The error bars denote the standard deviation. D. Accuracy of blastocyst prediction for maternally-young and aged embryos with and without developmental time indication when taken together with a selected set of morphological features.

To benchmark the effect of developmental times, we calculated the amount of information gained about survival given the embryo’s stage but without the particular transition times. In embryos from maternally old females, for example, knowing that the embryo has reached the early blastocyst stage carries more information about survival, compared to knowing it reached the 8-cell stage. We show that adding the knowledge of particular timing significantly improves the amount of information about survival (Fig. 2C). To test that developmental timing is not only informative, but has an explicit predictive power, we trained a machine-learning algorithm to predict whether an embryo would reach the late blastocyst stage using the eight significant features (Methods, Figure 2D). We show that adding the developmental timing as a bio-marker significantly improves the prediction accuracy, for both age groups and particularly for embryos from older females. For example, for embryos from older mothers at the morula stage, adding the developmental times improves the prediction, compared to using only morphological features, by around 10%.

Since timing is informative and contributes to prediction accuracy, we further investigated the distribution of developmental timing and its age dependency. A key question is whether embryos that reached the blastocyst stage have a different timing trajectory than embryos that did not. First, it is insightful to consider a few limiting scenarios for the interplay between timing, survival, and maternal age (Figure 3A). One possibility is an overall difference between embryos that survived or not, e.g., viable embryos are faster or slower than the non-viable ones (Figure 3A, top- left and middle panels).

Another option is that the difference between viable and non-viable embryos is the width of their temporal distribution, e.g., viable embryos are less ‘noisy’ (Figure 3A, top-right panel). The effect of maternal age can also result in similar limiting scenarios: older embryos are slower or faster than young ones, or have different temporal distribution width (Figure 3A, bottom panel). To answer these questions, we quantified the developmental timing distributions as a function of stage, viability, and maternal age (Figure 3).

**Figure 3.**
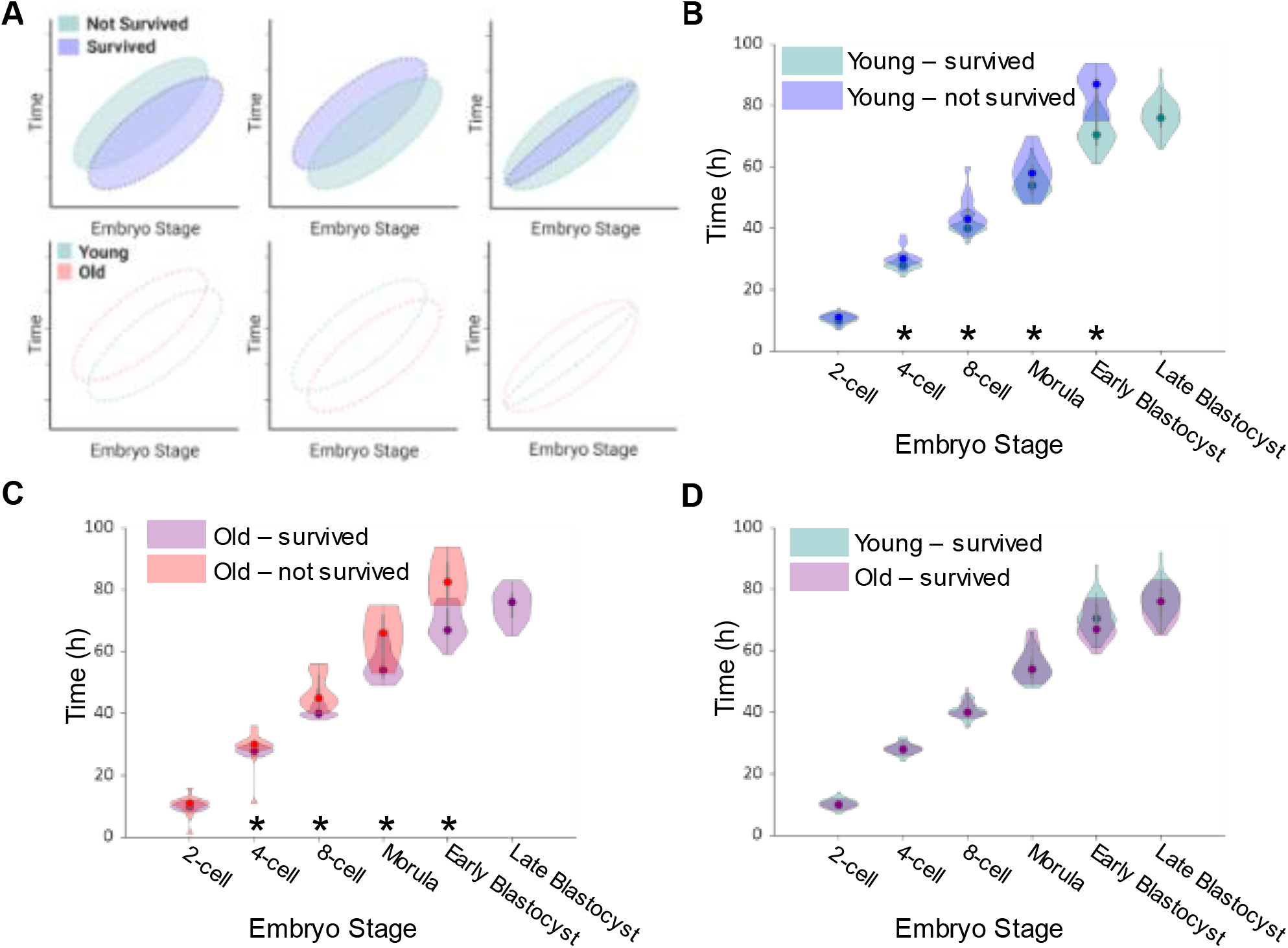
Embryos from both maternally aged and young females that reach the blastocysts stage follow the same tight developmental timing. A. Suggested optimal window for developmental time, independently of age (top). Suggested correlation between optimal developmental times for different age groups (bottom). B. Violin plots of development time for the various stages of embryos from young females. Embryos that do not reach the blastocyst stage tend to be slower. (*) denote differences that have a p-value ≤ 0.05, according to Kolmogorov-Smirnov test. C. For embryos from old females, the slowdown of embryos that do not reach the blastocyst stage is more pronounced. D. The timing of embryos that reach the blastocyst stage does not significantly depend on maternal age.

Our results show that maternally young embryos that fail to reach the blastocyst stage tend to develop significantly slower (Figure 3B). For maternally old embryos, the slowdown of the embryos that do not reach the blastocyst stage is even more significant (Figure 3C). Interestingly, the timing of the embryos that reach the blastocyst stage does not depend significantly on the mother’s age (Figure 3D). We show that viable embryos have a developmental timing window and that the hallmark of the non-viable embryos is a deviation from this window toward slower developmental timing. Altogether, our results suggest that viable embryos develop within a temporal corridor and that deviation from it is maternal age-dependent.

To further characterize the effect of maternal age on the transition timing, one needs to go beyond the distributions of transition times per stage and also account for temporal correlations. For example, would the embryo that entered first the 2-cell stage, will also be the first to enter the early blastocyst stage? Figure 4A illustrates the relation between transition times of consecutive stages. Almost all of the adjacent stage pairs exhibit significant correlation. Temporal correlation descends as the developmental window widens (see Figure 4A, inset), thus suggesting the existence of a memory that diminishes with further development. Interestingly, the developmental timing is independent of the embryo size (Figure S2).

**Figure 4.**
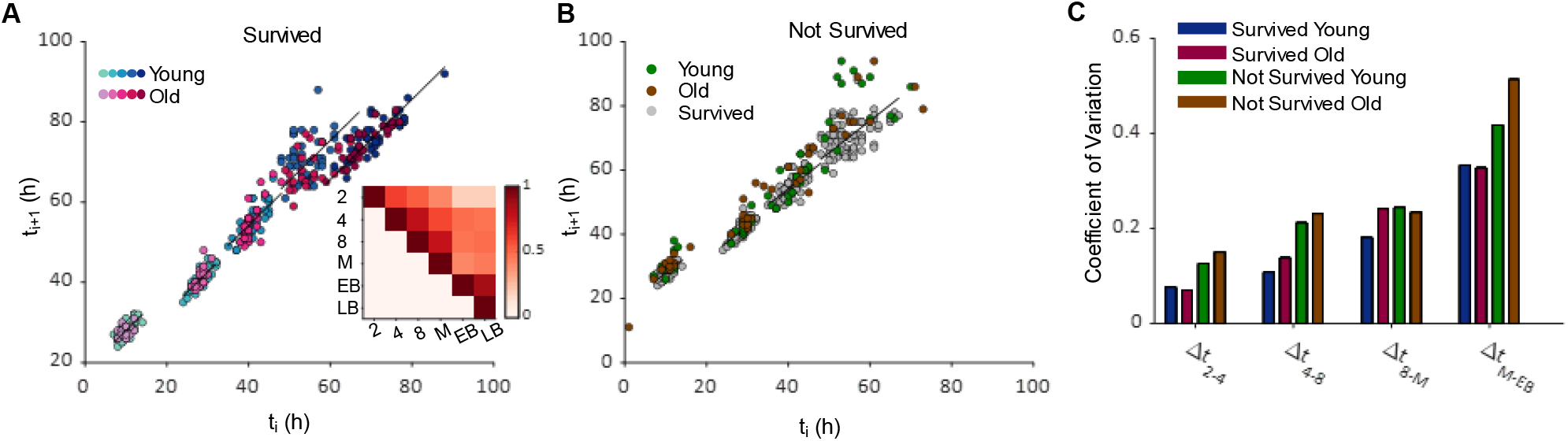
Temporal correlation between developmental stages is age dependent. A. Time correlation between adjacent stages for maternally-young (blue gradient) and old (red gradient) embryos that survived to the late blastocyst stage. Black lines are reference to perfect linear correlation with slope of 1. Inset heat map depicts correlation coefficient for developmental time for all survived embryos and stage combinations. M, EB, LB denote morula, early blastocyst and late blastocyst, respectively. B. Time correlation as in A, for maternally-young (green) and old (brown) embryos that didn’t survive to the late blastocyst stage. Surviving embryos are represented in grey for comparison. C. Bar plot for coefficient of variation as a function of time difference between each pair of adjacent stages. Maternal age and surviving state are represented by different group colors.

For embryos that did not survive up to the late blastocyst stage, the temporal distribution is noisier, especially for maternally-old embryos (Figure 4B). We have compared the coefficient of variation (CV) of temporal correlation between survived and non-survived embryos and different age groups (Figure 4C). One of the hallmarks of the non-viable embryo is the loss of correlation between consecutive stages. Moreover, this loss of correlation is maternal age-dependent. This is particularly evident at the transition from 8-cell to morula: embryos from old mothers exhibit a larger variation in their temporal difference than embryos from young ones.

## Discussion

Advances in assisted reproductive technology (ART) techniques provide a solution for many infertility problems and give rise to the increased use of IVF procedures worldwide. Almost a quarter of ART cycles in the US are done by women that are older than 40. The increasing age poses a significant challenge for ART success since both transfer and pregnancy rates decline with age^21,22^. To study the effect of maternal age on the pre-implantation mammalian embryo, we analyzed the development of mice embryos from the zygote to the late blastocyst stage. To quantitatively capture the entire morphokinetic properties, we have developed a machine learning platform that can not only track the cells and infer morphological properties but also determine their developmental stage in real-time.

First, we showed that, as expected, embryos from older females have a higher hazard risk (Figure 2A). Our aim was to quantify whether there are properties that are maternal-age dependent: Are embryos from older mothers tend to be bigger? Do they have different textures? Do they have different dynamics? These questions are also instrumental for ART where embryo grading is based on assessing the embryo form. To reveal how maternal age affects the developmental trajectory, we measured 45 morphological and textural features. In addition, our AI platform allowed us also to properly determine the transition times between developmental stages, a task that is laborious, time-consuming, and prone to errors.

We show that the property that is most informative regarding whether an embryo will reach the blastocyst stage is the transition time between developmental stages. This is true in particular for embryos from old mothers. The maximal information one can gain about whether an embryo is viable or not is one bit. Knowing the transition times provide 0.2-0.3 bit of information for embryos from old females (Figure 2B, 2C). Moreover, this knowledge translates into predictive power (Figure 2D). Knowing the transition times, improves the accuracy of predicting whether an embryo will reach the blastocyst stage or not. In particular, it has a significant impact on embryos from old females in their morula or early blastocyst stages. In the case of embryos at their early blastocyst stage, knowing the timing, brings the prediction accuracy of embryos from old females to that of embryos from young females.

To understand the interplay between maternal age, embryo viability and timing, we estimated the distribution of transition times at each stage. We found that non-viable embryos tend to be slower than the viable ones. Interestingly, we found that the distribution of transition times for viable embryos is maternal age-independent. However, embryos from maternally aged females, that are not viable tend to be even slower than the non-viable embryos from young mothers. Overall, our results suggest that embryos must remain within a tight temporal trajectory and that embryos from maternally aged females have a higher tendency to deviate from this corridor, and when they fail, they differ from it more than embryos from young mothers.

We also characterized the correlation between the transition times of the same embryo. We show that transition times between consecutive stages are significantly correlated (Figure 4A). Moreover, the correlation decreases when comparing stages that are further away from each other but is not fully diminished, thus suggesting some memory along the developmental trajectory. The temporal correlation of maternally aged embryos tends to be lower and the coefficient of variation in the difference between transition times is higher (Figure 4C). These results also suggest that the development timing is under a tight regulation and that embryos from older mothers have a higher tendency to delay their developmental transition. Interestingly, the transition times show no dependence on the size of the embryo (Figure S2), that is, small embryo or large embryo have the same timing. Our results suggest that viability is associated with a tight timing and that embryos that are too slow tend to not reach the blastocyst stage. This emerging picture of the embryo as a timer is consistent with the need for temporal coordination with the uterus that needs to be ready to promote proper implantation. Our results highlight the importance of developmental transition times, particularly for embryos from old females, and pave the way for increasing ART success rate in older women.

## STAR Methods

### KEY RESOURCES TABLE

| REAGENT or RESOURCE | SOURCE | IDENTIFIER |
| --- | --- | --- |
| Software and algorithms |  |  |
| MATLAB R2018a, version 9.4.0.813654 | The Mathworks | SCR_001622 |

### RESOURCE AVAILABILITY

#### Lead contact

Further information and requests for resources should be directed to the lead contact, Yonatan Savir.

#### Materials availability

There is no restriction for distribution of materials.

### METHOD DETAILS

#### Ethics Statement

All mouse experiments performed in this study were approved by the Animal Care and Use Committee of the Technion, Israel Institute of Technology, and found to conform with the regulations of this Institution for work with laboratory animals.

#### Collection of Mouse Embryos and Live Imaging

Embryos were collected from F1 (C57BL/6×DBA) females (Young n=5, Old n=4) induced to superovulate by intraperitoneal injection of 7.5 IU of pregnant mare serum gonadotrophin (PMSG) (Sigma Aldrich) followed 48 hours later by 7.5 IU of human chorionic gonadotrophin (hCG) (Sigma Aldrich) and then mated with males of the same genotype. Mice were euthanized by cervical dislocation and eggs were collected 16.5 hours after hCG injection into KSOM (Merck) media containing 200 IU/ml of hyaluronidase (Sigma Aldrich), dispersed and then transferred to fresh KSOM media. Only morphologically competent eggs were selected for culturing. Embryos were cultured in 5% CO2 at 37°C in a humidified incubator and observed under an inverted (Nikon) microscope using DIC optics and 20X/0.75 NA objective. DIC image was acquired every 60 min, with an exposure of 10 ms for transmitted light for total of 96 hours. A total of 103 embryos (Young n=62, Old n=41) were further implemented in our experiment.

#### Network Training

We labeled about 3000 images of the embryos in different stages (from zygote to blastocyst). We randomly divided the categories into 70% images for training and 30% for testing. We used a variety of residual and convolutional networks (Fig. S1). During the training, different hyperparameters were examined. The initial learning rate was 0.0001 with a drop factor of 0.5, and a period factor of 5 (piecewise schedule). The net optimizer was Adam, with an input size of 224X224 pixels, mini-batch size of 24, decay factor of 0.99, 10 epochs, training iteration number of 148, and testing iteration number of 50.

#### Morphological Features

45 morphological textural and geometric features were extracted for each embryo as a function of time. In particular, we used standard geometric features: Area, Diameter, Perimeter, Roundness, and Eccentricity. We also included a set of 20 well-known textural features extracted from the Gray-Level Co-occurrence matrix^23,24^: Autocorrelation, Contrast, Correlation, Cluster Prominence, Cluster Shade, Dissimilarity, Energy, Entropy, Homogeneity, Maximum Probability, Variance, Smoothness, Sum Average, Sum Variance, Sum Entropy, Difference Variance, Difference Entropy, Info Measure of Correlation, Inverse Difference Normalized, Inverse Difference Moment. A set of textural features from Histogram of Oriented Gradients (HOG) also included^25–27^ a mean value of HOG descriptor. A set of statistical textural features from Gray-Level Difference method^28^: Horizontal Weighted Sum (HWS), Vertical Weighted Sum (VWS), Diagonal Weighted Sum (DWS), and Grid Weighted Sum (GWS). Image gradient-based features, which are standard functions for extracting homogeneous patterns, boundaries, edges of the image: gradient magnitude, and gradient direction values. Textural features of Total Variation (TV) transform ^29^: Low Pass Spectral Filtering TV, High Pass Spectral Filtering TV, and Band Pass Spectral Filtering TV. Co-occurrence probabilities features extracted from embryo images after High Pass Spectral Filtering based on TV-transform^23,24,29^: Texture Skewness, Gradient Mean, Gradient STD, Gradient Global Mean, Gradient Uniformity, Gradient Entropy, Gradient Skewness, Gradient Correlation.

#### Mutual Information

The mutual information is given by 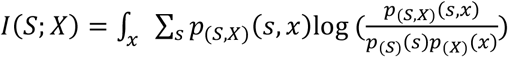, where *s* is the indicator variable for reaching the blastocyst stage, and *x* is a feature. To account for sampling size, we bootstrapped the mutual information for each stage, for each feature, for each age, 100 times, assuming x is drawn randomly from the observed distribution that leads to a distribution of basal information, *I*, values. Only features that provided an information gain that is one standard deviation above the mean basal *I* are shown in figure 2B.

#### Predicting Blastocyst Formation Rate

We used Matlab to train 3 different Machine Learning methods: SVM, LD, and ensemble models. To predict the blastocyst survival rates based on the informative features (Figure 2B): Timing, Cluster Prominence, Cluster Shade, Smoothness, Gradient Uniformity, Gradient Correlation, Low Pass Spectral Filtering TV, and standard deviation of the gradient magnitude of the segmented embryo (V_G_mag). Ensemble model using subspace method, discriminant learner, 30 cycles gave the best result.

