## Supplementary Information for "Machine Learning Reveals The Effect of Maternal Age on The Mouse Pre-Implantation Embryo Developmental Timing"

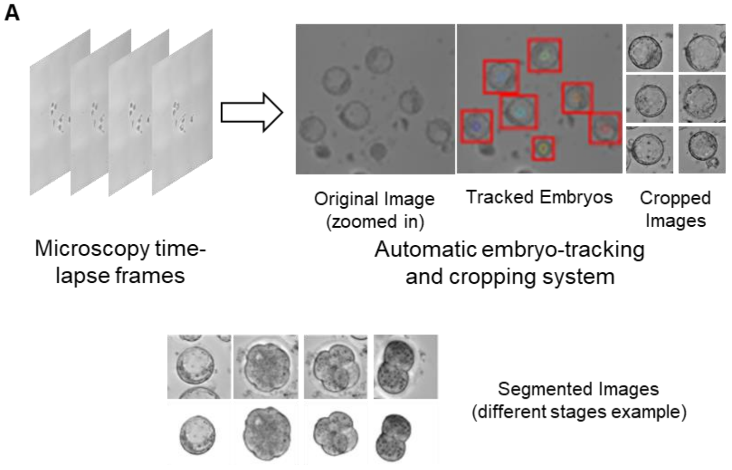


B

| Model | % Accuracy |
|---|---|
| VGG16 | 92.21% |
| ResNet-18 | 87.37% |
| ResNet-50 | 97.17% |
| Inception-v3 | 89.23% |
| AlexNet | 86.90% |
| GoogleNet | 96.20% |

C

| Stage | 2-cell | 4-cell | 8-cell | Morula | EB | LB |
|---|---|---|---|---|---|---|
| Mean Timing Error(hr) | 0.0291 | 0.2451 | 0.3069 | 0.52 | 0.4556 | 2.1111 |
| STD of Timing Error | 0.1690 | 0.6204 | 0.6123 | 1.8775 | 1.0722 | 2.7962 |

**Figure S1.** Artificial intelligence pipeline structure and accuracy. A. Deep learning platform pipeline. Automated tracking and cropping of the embryos are performed, after which the image set is used for training. B. Comparison of accuracy between different network architectures. C. Mean timing error for each developmental stage and error standard deviation in hours. EB denotes early blastocyst. LB denotes late blastocyst.

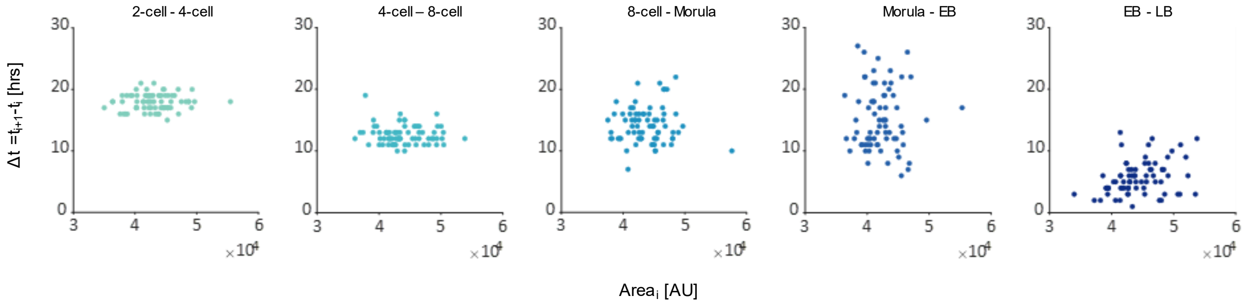


**Figure S2.** Developmental time is size-independent. A scatter plot of Δt between each pair of consecutive stages as a function of the initial area. EB denotes early blastocyst. LB denotes late blastocyst.
